# Exposure of non-target white-footed mice (*Peromyscus leucopus*) to Second-generation Anticoagulant Rodenticides in an urban context

**DOI:** 10.64898/2026.04.06.715702

**Authors:** Luigi F. Richardson, Albrecht Schulte-Hostedde

## Abstract

The pathways of non-target exposure to anticoagulant rodenticides (ARs) are poorly understood, and have yet to be examined in Ontario, Canada. The spillover of ARs into non-target rodents and high-risk landscapes has been investigated numerous times, but usually in agricultural regions as opposed to urban ones. We used snap traps to capture white-footed mice (*Peromyscus leucopus*) in urban wildland areas of Toronto and Vaughan, Ontario near ongoing rodenticide baiting programs. Our goal was to determine if second-generation anticoagulant (SGAR) baiting practices used by pest management professionals targeting commensal rodents may be causing rodenticide spillover into non-target rodents in urban wildland areas, which could act as a vector of ARs to predators. Only 11 out of 111 mature white-footed mice trapped near ongoing urban rodenticide operations tested positive for an anticoagulant, at five out of seven study sites. Concentrations were between 0.008-0.03 ppm, which may be sublethal for raptors. We did not detect brodifacoum, despite its detection in a recent study on Ontario raptors. Exposed individuals were caught at 0m, 5m, 20m, 40m, 70m and 100m from active rodenticide stations. They did not differ from unexposed individuals in terms of sex, age, body condition, distance to the AR source, capture date or capture site. This indicates that the pest management industry’s use of rodenticides in urban and suburban settings is causing some degree of non-target spillover in Toronto and the Greater Toronto Area, and that SGAR usage should be avoided near naturalized landscapes.

## Introduction

Anticoagulant rodenticides (ARs) are the gold standard for low-maintenance rodent control programs worldwide (Endepols 2002). ARs rose to popularity because of their ability to prevent bait shyness through delayed effect lethality. Predators that consume rodents during the latent period before death can thus be exposed to ARs (Sainsbury et al. 2018). Second-generation anticoagulants (SGARs) are currently the most popular variety of ARs, and are known for their propensity to bioaccumulate, and magnify in higher trophic levels (Jaramillo-Q. et al. 2024). The exceedingly toxic SGARs were designed specifically to control “warfarin-resistant” rodents (Boyle 1960), which evolved insusceptibility to the first-generation of these poisons (Rost et al. 2004). However, the exact pathways of non-target exposure are not well understood (Elmeros et al. 2019), making the creation of responsible rodenticide usage programs particularly difficult. This is a particular issue of concern in Canada, where research on rodenticide resistance, predator exposure and prevention is sparse (Thomas et al. 2011; Elliott et al. 2014; Thornton et al. 2022).

Peridomestic rodent species use human habitations and structures intermittently (Douglass et al. 2006), unlike commensals, which are characteristically synanthropic and reliant on human activity (Weissbrod et al. 2017). Peridomestics may be the main focus of rodent control programs or end up as bycatch, and they may also occupy different spatial niches where they are sympatric with commensals (Frye et al. 2021). Given that the majority of pest management professionals (PMPs) in Canada have few commercial first generation anticoagulant (FGAR) alternatives for rodent infestations, it is likely that peridomestic, native non-target rodents (NTR), like *Peromyscus spp*. (deer and white-footed mice) are heavily exposed to super-toxic SGARs, as is the case with non-targets in other regions (Elmeros et al. 2019). They can move great distances from infested buildings quickly (Calisher et al. 1999), and may have a preference for natural landscapes (Hummell et al. 2023) where predators may be more abundant, making them a main pathway of secondary exposure (Geduhn et al. 2016). They may also be the preferred prey of Ontario’s predators (Baker and Brooks 1981). Commensals are the sole target of SGARs in Canada, and are often less available to predators after poisoning because of their stricter synanthropy; they are often subterranean, and more abundant inside and around human structures where predators may be less prevalent (Kaufman and Kaufman 1990; Geduhn et al. 2014; Walther et al. 2021). In addition, although not registered or designed for the control of native NTR, PMPs can still perform general rodent control outside and around structures and fence lines using SGARs (i.e. bromadiolone) in Canada (Health Canada 2012), possibly exposing *Peromyscus spp*. or other non-targets to the chemical (Hindmarch et al. 2018). PMPs and farmers are the only parties authorized to apply SGARs in Canada (Health Canada 2012). Most studies of primary and secondary non-target exposure focus on agricultural use patterns (Tosh et al. 2012), but which of these two parties may be more responsible for AR spillover is unknown.

Scientists often choose to study the fate of poisoned rodents themselves rather than testing submitted carcasses to approximate the secondary poisoning risk to non-targets in different contexts (Hughes et al. 2013), in order to provide a less biased gauge of risk associated with AR usage practices (Tosh et al. 2011). Variation in exposure risk can be affected by the rodent species being exposed, which may be related to landscape factors, as well as the distance to the AR source (Geduhn et al. 2014, 2016). Males and mature rodents may be exposed at higher rates or to greater concentrations of ARs as a result of their larger home range (Metzgar 1973; Wolff 1985; Gaitan and Millien 2016), which may change seasonally (Getz 1961). Greater quantities of rodenticide at an extermination site may correspond with higher exposure risk, especially among rodents found closer to bait stations (Tosh et al. 2012). Non-target *Peromyscus spp*. mice may enter structures (Douglass et al. 2006; Frye et al. 2021), and SGARs may be deployed before visual species identification occurs (Hindmarch et al. 2018). It is also well known that *Peromyscus sp*. will enter rodenticide stations alongside commensals (Baldwin et al. 2014).

Permanent rodenticide baiting, such as that which occurs in Canada on commercial properties or in agricultural areas (Hindmarch et al. 2018) is cited as a principal cause for non-target rodent and predator exposure to ARs in Europe, particularly in agricultural settings, likely as a result of the exposure of similar non-targets like *Apodemus spp*. (field and wood mice, Endepols et al. 2015; Geduhn et al. 2016; Elmeros et al. 2019). These NTR are known to subsequently retreat to open fields or wildland areas, thereby acting as a “vector” through which rodenticides enter predator-friendly landscapes (Tosh et al. 2012). Various commercial and food handling establishments often do or are required to employ professional pest management services in Ontario under regulation 493/17 (Government of Ontario 2017), and these programs commonly utilize some form of rodenticide treatment for the duration of the contract. The percentages of sampled NTR exposed to rodenticides varies in studies of secondary poisoning pathways but is typically between roughly 12-48% (Endepols et al. 2003; Brakes and Smith 2005; Walther et al. 2021). No exact rate of exposure appears to be associated with hazard levels that put predator species’ populations at risk, but patterns related to increased exposure rates can be used to correct SGAR usage programs (CRRU 2018).

We sampled white-footed mice *Peromyscus leucopus* from fragmented urban forests adjacent to commercial buildings using permanent SGAR baiting to determine if this non-target species is being exposed to SGARs, and if exposure varied with sex, age, season, amount of bait used and distance from the SGAR source. We hypothesized that non-target white-footed mice would be exposed to SGARs as a function of their sex, age class and distance from rodenticide bait stations in urban landscapes. We predicted that males and mature individuals would be exposed more often and to higher concentrations of AR chemicals, especially when closer to exterior bait stations, as a result of their larger home range (Metzgar 1973; Wolff 1985; Gaitan and Millien 2016), and that bromadiolone would be the most commonly-detected chemical in white-footed mouse liver tissue, due to its popularity (Elliott et al. 2014) and availability for outdoor usage (Health Canada 2012).

## Methods

### Field Methods

All trapping was conducted under an Ontario Ministry of Natural Resources and Forestry (OMNRF) Wildlife scientific collector’s permit (WSCA 1106092), and the approval of Laurentian University’s Animal Care Committee. White-footed mice were captured by snap trapping (lethal sampling) using Victor® M7 snap traps inside secured bait stations (Bell Laboratories Protecta EVO^®^ EXPRESS™). Stations were modified with sheet metal to prevent the entry of larger animals through the ingress point (similar to Bell et al. 2019). Similar trapping methods have been used previously for gauging the degree to which small mammals available to predators may be contaminated by rodenticide residues (Brakes and Smith 2005; Sage et al. 2008; Lima and Salmon 2010; Winters et al. 2010; Tosh et al. 2011; Elliott et al. 2014; Geduhn et al. 2016; Desvars-Larrive et al. 2017; Elmeros et al. 2019; Murray and Sánchez 2021; Walther et al. 2021a; Mahamat et al. 2024).

Seven “wildland” sites (municipal parks composed of mixed deciduous forests) across Toronto and Vaughan, Ontario, adjacent to commercial buildings with ongoing, permanent exterior rodenticide programs, were selected as sites for trapping. Rodenticide program details were confirmed through phone call discussions or emails with property managers. The general days or weeks when bait stations were refilled (e.g. every first and third week of the month) were provided also and were used to approximate the elapsed time from rodenticide applications relative to sampling days. Sites one, two, three and four were in Vaughan, site five was in Scarborough, site six was in North York and site seven was in Etobicoke (see supplementary material). Five sampling sites were owned by the Toronto and Region Conservation Authority and two were owned by the City of Vaughan. Written permission for trapping was obtained from both parties before the sampling period began on June 28^th^, 2024. Sites were trapped once every three weeks until the end of August, to obtain rodent carcasses at different times in relation to scheduled PMP site visits, when rodenticide refills may occur. A total of three trapping events, when traps were set and checked the following day, occurred per site, and overall there were 334 trap nights.

At each site, a transect was created as the basis for a trapline using ArcGIS^®^ Online. The transect was drawn perpendicular to the foundation wall of the building where AR was in use, and trapping stations were placed at specific, uniform distances (see supplementary material). Coordinates were stored on a Garmin® GPS map 64s and used for setting stations in the field. Distances chosen were 20 metres, 40 metres, 70 metres and 100 metres, approximating the distance intervals used in previous studies (Elliott et al. 2014; Elmeros et al. 2019). The width of chosen wildland areas restricted our ability to have each of these exact distance points at each site. The 20-metre distance was used for sites two, three, six and seven; 40m and 70m for all sites; 100m for sites one, two, four, five and six. Trap stations were also placed at the edge of the wildland area (either five, 15, 20 or 40 metres from the base of the foundation). In one case, the edge of a wildland was 31 metres from the rodenticide use building but was still incorporated into the 40m category for analytical purposes. Finally, to quantify urban lands within each trapping location on the transect line, the percentage of developed space (buildings, hardscapes and manicured lawns) was assessed at each distance location within the estimated home range size of *P. leucopus* (7,600 m^2^ circular area, Gaitan and Millien 2016).

Two trapping stations were placed within two metres of each coordinate location, with entry holes facing north-south, and labelled “A” or “B”. By extension, traps inside were labelled N or S (north/south). Stations were left in place with unset traps baited with peanut butter and hamster feed for between one and six weeks. Traps were set on weekends only (Fridays or Saturdays) and checked the following day within 24 hours (Saturday/Sunday), and left unset until the next trapping period. Trapped rodents were removed, and sex, age, body weight and length were recorded. Carcasses were placed in labelled storage bags detailing location and frozen on the day of capture. Only mature adult white-footed mice, with testes present or perforated vaginas, were used for liver analysis. Liver samples were sent to the University of Guelph Animal Health Lab for liquid chromatography tandem mass spectrometry testing (methods similar to Thornton et al. 2022). Anticoagulant rodenticide analytes included: brodifacoum, bromadiolone, difethialone, flocoumafen, difenacoum, warfarin, chlorophacinone, diphacinone, pindone, dicoumarol, coumatetralyl, coumachlor, coumafuryl and valone.

### Statistical Methods

Contingency tables were used to conduct Fisher’s tests to compare frequencies of exposed and unexposed white-footed mice based on sex, age or capture site. Length (millimetres) and weight (grams) were graphed using linear regression to calculate body weight residual as a metric of body condition, as a function of the individual’s distance from the least squares line of best fit (i.e. negative or positive residual). This value was then used as a predictor of exposure. Non-parametric Mann-Witney U tests were used to compare body morphometry measurements and environmental factors that may relate to exposure at the capture site. Factors used for predicting rodenticide exposure and concentration included rodent sex, rodent age, the number of rodenticide stations at the adjacent building, percentage of developed land within the estimated home range, distance from AR source, and Julian date of capture. We compared these metrics between exposed (AR positive) and unexposed (AR negative) individuals. Effect sizes were also calculated. Linear regression was also used to compare these variables to concentration of AR chemical detected, and checked for heteroskedasticity and normality of residuals with Breusch-Pagan and Shapiro-Wilk tests, respectively. Principal component analysis was then used to capture covariation between these variables, and regression was performed with PC1 against the AR concentrations of exposed individuals. Logistic regression was used to determine if the number of bait stations at a site had any relationship with exposure status (exposed or unexposed). Statistical analyses were performed in R Studio (R Core Team 2024, R version 4.4.2). Packages used included “lmtest” and “RStatix”.

## Ethics Statement

All trapping collections were conducted under an Ontario Ministry of Natural Resources and Forestry (OMNRF) Wildlife scientific collector’s permit (WSCA 1106092), and the approval of Laurentian University’s Animal Care Committee. Open AI’s ChatGPT was used to assist in writing R studio scripts for data analysis, which were subsequently checked for correctness and modified for customization.

## Results

Trapping success for all seven sites was 47% (158 captures/334 trap nights). A total of 11/111 adult white-footed mice tested were positive for anticoagulant rodenticides (9.9%, see Table 1), all of which were second-generation compounds (SGARs); either difethialone (two) or bromadiolone (nine). Exposed mice were caught at sites one, three and four (Vaughan), as well as six and seven (Toronto) (see supplementary material).

**Table 1.** Comparison of parameters for rodenticide-exposed (Pos.) & unexposed (Neg.) individual white-footed mice. The following parameters were compared between exposed and unexposed individuals: a) Body length (mm) b) Weight (g) c) Body weight residual (g) and d) % Urban land within estimated home range at the capture site.

|  | AR Positive (n = 11) |  | AR Negative (n = 100) |  | MW *p* value | Effect size (*r*) |
| --- | --- | --- | --- | --- | --- | --- |
|  | Median | (IQR) | Median | (IQR) |  |  |
| Body length (mm) | 94 | (12.5) | 90 | (8) | 0.82 | 0.02 |
| Weight (g) | 25 | (7) | 21 | (5) | 0.22 | 0.12 |
| Body weight residual (g) | 0.48 | (3.37) | 0.07 | (2.89) | 0.11 | 0.15 |
| % Urban land | 15.79% | (38.55%) | 13.71% | (39.62%) | 0.29 | 0.08 |
| *Fisher's exact test for exposure rate of mature and immature males and females = 0.45* |  |  |  |  |  |  |

Concentrations of bromadiolone ranged between 0.008-0.03 ug/g of liver tissue (av. 0.015 ug/g) and were 0.008 and 0.023 ug/g for difethialone. Difethialone was only found at sites three and four (both in Vaughan, separated by Rutherford Road by < 100m). Never was more than one SGAR detected in a single individual mouse. All sites were mixed deciduous-coniferous forests, and there was no difference in the proportion of exposed individuals caught at these five sites (fisher’s exact p = 0.39). Two or more exposed individuals were never caught at the same distance at a single site, meaning that a single trap box or trap never caught more than one exposed mouse. One exposed *P. leucopus* was caught 100m from a rodenticide use building, and one was caught directly beside a bait station on one occasion (0m).

Of the exposed individuals, two were immature females (18%), two were adult females (18%) and 7 were adult males (64%). There was no significant difference in the proportion of exposed vs unexposed males and females (fisher’s exact p = 1) or matures and immatures (fisher’s exact p = 0.68). Mann-Witney U tests were used for unifactorial comparisons of exposed (n = 11) and non-exposed (n= 100) individuals, and there was no difference between these groups in terms of body length, weight, body condition, or the percentage of urban land at the capture location (all p > 0.05, Table 1). Despite the lack of statistically significant differences, exposed white-footed mice were, based on medians, longer, heavier, in better condition, and caught at locations with a greater percentage of developed land within their estimated home range (Table 1). Two individuals (18%) were caught within one week of a scheduled PMP site visit, five (45%) were caught within two weeks, two (18%) were caught within three weeks, and two (18%) were caught within four weeks (these were the only four categories).

Predictors were incorporated for PCA (Urban land %, sex, age class, Julian Date of Capture, distance from AR building), and there was no significant correlation between exposure concentration of bromadiolone and principal component one (p = 0.15). Difethialone-exposed mice (n = 2) were not incorporated in this analysis, because this SGAR is formulated at 0.0025%, as opposed to 0.005%, in rodenticide baits (Health Canada 2019). PC1 explained 66.04% of the variance across *%UrbanLand* (0.91), *Sex* (0.55), *Age* (-0.95), *Julian date* (-0.85) and *Distance* (-0.72). There was also no correlation between exposure concentration or any of the predictors singly (all p > 0.05). Finally, the number of bait stations at each site was not related to AR exposure status of mice (logistic regression p = 0.16, see supplementary material).

## Discussion

We attempted to determine factors associated with SGAR exposure in a non-target urban rodent species (*Peromyscus leucopus)* resulting from ongoing commercial SGAR usage, in Toronto and Vaughan, Ontario. This is one of the few observational, *in-situ* examinations of real-world AR spillover from urban commercial operations, as opposed to agricultural studies (Geduhn et al. 2016) and controlled studies where researchers simulate AR usage by placing their own bait stations (Elmeros et al. 2019), or through modelling (Topping and Elmeros 2016). Although *Peromyscus spp*. are known to use bait stations set for the control of commensals (Burke et al. 2021), wild-caught *Peromyscus spp*. appear to have only been previously tested for rodenticide residues twice: Elliott et al. (2014) found no residues in three *P. maniculatus* total, and Lima and Salmon (2010) found trace levels (sample size not given). Our data show no significant correlates or predictors for exposure, or differences between exposed and unexposed mice.

Our overall exposure prevalence was relatively low (9.9%, 11/111), and the most commonly detected chemical was bromadiolone (9/11). Comparable studies using rodent liver tissue as an AR detection medium have typically found exposure rates between 12-48% in peridomestic and non-target rodents, namely, *Apodemus spp*., *Clethrionomys spp*., *Microtus spp*. and *Arvicola spp*. (Brakes and Smith 2005; Sage et al. 2008; Tosh et al. 2011; Geduhn et al. 2014; Elmeros et al. 2019). However, these studies were all conducted in agricultural settings, which have less contrast with surrounding natural landscapes than urban areas which may deter rodent travel and foraging (Ortiz□Jimenez et al. 2025). NTR may therefore be more receptive to rodenticide bait stations in agricultural regions (Patergnani et al. 2010). Additionally, despite liver tissue typically holding the majority of AR residues (Rached et al. 2022), using homogenized whole-body tissue may have yielded much higher rates of exposure (Sage et al. 2008). Our effect sizes for unifactorial comparisons were all well below the threshold even for small effects (Table 1), indicating that exceedingly large sample sizes would be needed to show significant differences between body morphometrics and urban land within estimated home ranges of exposed and unexposed mice.

A previous study on raptors collected throughout the province of Ontario also showed bromadiolone to be the most prevalent chemical detected, followed by brodifacoum (which we did not detect) and difethialone, but at much greater concentrations and prevalence rates than we present here (Thornton et al. 2022). Considering the biomagnifying nature of SGARs (López-Perea et al. 2019; Buckley et al. 2024) and the trophic level of raptors relative to *Peromyscus spp*., this is axiomatic. However, in contrast with our results, it may also indicate that raptors could be a vacuum for AR poisoned rodents, selectively choosing them (Cox and Smith 1990), or at least, consuming more proportionally than a trapping study could ever detect.

Bromadiolone was detected from *P. leucopus* caught in close proximity (0-20m) and further away (40, 70 and 100m) from active bait stations, with an average concentration of 0.015 ug/g. Apparently, no LD50 values are available for *Peromyscus leucopus*., but considering previous studies have shown 100% susceptibility of *P. maniculatus* to 0.02% (200ppm) chlorophacinone after three days feeding (Marsh et al. 1977), the concentrations we detected may have been lethal, and are comparable to those found for NTR in previous studies (Geduhn et al. 2014). These doses could affect a great number of wildlife that may be secondarily exposed, including raptors, considering different taxa are variable in their susceptibility to ARs (Lemus et al. 2011; Eckel et al. 2020), and residues may persist for some time after the exposure in an otherwise healthy animal. However, thresholds for sublethal and lethal effects are reportedly between 0.1-0.2 ug/g; far higher than any of our detected concentrations (Carromeu-Santos et al. 2026). We also found no significant correlation between distance from the source and concentration of AR chemical in liver tissue. However, regarding brodifacoum (formulated at either 50 or 25 ppm in Canada), liver residues rarely exceed 1 ug/g, and the highest concentrations are typically found during the AR baiting period (Geduhn et al. 2014). Concentrations are also typically higher in target species than non-targets (Geduhn et al. 2014; Walther et al. 2021) and at closer distances to the source (Elmeros et al. 2019), but not always (Tosh et al. 2012).

The fact that our trapping success, a proxy for rodent density (Kaboodvandpour et al. 2010), was relatively high at all sites (47% catch rate/334 trap nights) suggests that rodenticide programs at the adjacent usage buildings may not be decreasing native rodent population density, and thus, the chemicals used may not spilling over enough to threaten predator populations. In contrast, prior studies showed that Prairie dog *Cynomys ludovicianus* control with ARs didn’t affect *Peromyscus* density (Deisch et al. 1990; McCaffrey et al. 2009), and others have shown that similar species’ populations (*Apodemus spp*.) can recover quickly from declines resulting from mass poisoning (Bell et al. 2019), meaning that white-footed mouse populations may persist despite anticoagulant pressure. Seasonal feeding and movement patterns may change this, however, and given that our trapping only took place in the summer, it may be possible that exposure rates, rodent density and associated secondary exposure risk could change as a function of seasonal factors (Colvin 1984; Tosh et al. 2011; Geduhn et al. 2016). Commercial facilities have previously noted that exterior AR stations may go untouched by target and non-target species alike for long periods of time, possibly due to the absence of rodents, spatial or temporal rodent feeding preferences (Hindmarch et al. 2018), and this, along with rodent species assemblages (Geduhn et al. 2016) may even vary with the seasons.

Rodents are known to remove and cache rodenticide baits (Brakes and Smith 2005), which could also explain our lower exposure prevalence rates, as they may continue to feed on them at a steady rate after their removal from the bait station, or abandon them. This could give the impression of lower exposure initially, but may result in exposure even after baits have been removed (Brakes and Smith 2005). Rodents may not consume this cached rodenticide for some time, and hence, depending on changes in food availability, exposure rates may rise when cached food provides the most attainable form of food (Brakes and Smith 2005). However, given that the risk of secondary exposure is said to be high in heterogeneous urban-wildland landscapes (Lohr 2018; Burke et al. 2021; Silveira et al. 2024), our low exposure rate may reflect that snap trapping is not a representative sampling method. Considering that lethal snap traps have previously been used to sample rodents for AR assays, and yielded higher rates of exposure (Geduhn et al. 2014), and the usefulness of snap traps in studies of rodent density and disease (Peitz et al. 2001; Karell et al. 2010; Marquez et al. 2019; Herawati and Purnawan 2021), this seems unlikely. Our low exposure rate could also indicate that patterns of urban exposure are different in Ontario than in other regions, possibly due to SGAR restrictions or use practices (Eckel et al. 2020), or that, for some other reason, we were unable to catch many exposed rodents.

Brodifacoum may not be favoured by industry professionals and may be used less often used than bromadiolone in Canada (Elliott et al. 2014), which may explain its absence from our samples. However, brodifacoum is typically the most frequently detected chemical in non-target rodents, corresponding with its half-life being greater than any AR available in many regions (Eckel et al. 2020), including Ontario (Thomas et al. 2017), but this may be due more so to its greater popularity and more frequent usage in other regions. Few baits available in Canada contain brodifacoum, none of which are in soft bait or liquid formulation (which may increase palatability to rodents), and these can only be applied indoors (Health Canada 2012). This may explain why brodifacoum was not detected in our samples, and why difethialone was only detected twice, at two locations within one hectare of each other. However, the frequent detection of brodifacoum in Ontario raptors (35%, Thornton et al. 2022) suggests that this exposure may be occurring by means of a different landscape or species exposure pathway.

Despite being an “indoor-only” chemical (Health Canada 2012), difethialone was detected 100m from a rodenticide use building at site four but could also have originated from residential usage at a nearby townhome complex (Bartos et al. 2011), or illegal outdoor usage (Thompson et al. 2014). In contrast, bromadiolone, which has likely been the most popular SGAR among Canadian commercial operators since the late 1990s (Elliott et al. 2014) is available in numerous block, pellet and soft bait products for indoor and outdoor usage. It was detected much more frequently in our samples (9/11 positives), though still at a low rate. Our detection of difethialone at two sites (three and four) indicates that white-footed mice may move indoors, consume SGAR chemicals which they should not be exposed to, and then travel some distance from the source before death, posing a risk to predators, assuming that difethialone was applied legally at these sites. The fact that no FGAR chemicals were detected, and that no exposed individuals tested positive for multiple chemicals, indicates that homeowners and commercial operators are not using such products, *P. leucopus* is not receptive to them or exposed to them, or that they are perhaps able to metabolize them (Poche and Mach 2000) below the detection levels of the equipment used. It may also be the case that a larger sample size would have allowed us to detect FGARs.

A recent study conducted in California determined that commensal rodents may be a main vehicle of Coyote *Canis latrans* exposure detected in the state, in spite of heavy AR restrictions (Stapp et al. 2025). However, given warmer year-round climates in the American west, the preponderance of black rats *Rattus rattus* in California, and the fact that we detected only one commensal at any of our trapping sites (1/254 total captures) suggests commensals may be able to uniquely capitalize on the California climate. They may become abundant in areas frequented by predators, and act as a vector of rodenticides (Stapp et al. 2025). The only commensal rodent we captured (house mouse) was trapped near a fenceline in a park, adjacent to a building where rodenticide usage could not be confirmed, for which reason this sample was not tested. Alternatively, the parking lots separating wildland areas from bait stations at our sites may have acted as an anthropogenic landscape of fear (Krijger et al. 2017), confining most *P. leucopus* to the wildland areas, which is a common occurrence in urban regions (Harris and Munshi□South 2017). Given that no exposure was detected at site five, where there was a unique rise in elevation and a drop of over 100m down a prominent slope, elevation may have some influence on the movement of rodents between wildland and urban areas, perhaps acting as a barrier against exposure. However, some degree of elevation drop was present at all other sites.

The fact that exposed rodents were caught at every distance location from identified rodenticide stations suggests that the likelihood of exposure may be more influenced by season, sex/age demographic or body morphology than proximity to the chemical source. Given that only 4/11 individuals were in “poor” body condition, larger and more dominant individuals, especially matures, may have better access to all local food sources than smaller subordinates. These larger prey species may also be preferred by raptors (Karell et al. 2010), and more likely to consume rodenticides (Caut et al. 2007). Most exposed individuals (7/11) were also caught within one or two weeks of scheduled PMP visits. This is consistent with previous research that suggests exposure rates climb shortly after baiting begins and decrease as the baiting period goes on (Walther et al. 2021). However, residues in dead rodents may be exceedingly high (Winters et al. 2010), meaning that any individuals which we may have missed using our trapping methods as a result of their being moribund or dead could represent the greatest, unknown risk. No dead rodents were ever found while setting up, checking or removing trapping stations in and around the transects.

## Acknowledgements

The authors would like to thank Dr. Claire Jardine, Dr. Catherine Cullingham and Dr. Jason Munshi-South for their initial reviews of the thesis from which this manuscript was derived. Also, the City of Toronto and City of Vaughan for allowing for the collection of rodents on municipal lands.

## Funding

This research was funded by an NSERC Discovery grant to AISH.

## Competing Interests

Luigi Richardson is and has been, since 2020, a pest management technician operating in Toronto and the greater Toronto area, with different companies.

## Author Contributions

Both authors contributed to the study conception and design. Material preparation, data collection and analysis were performed by L. Richardson. The first draft of the manuscript was written by L. Richardson and A. Schulte-Hostedde edited previous versions of the manuscript. Both authors read and approved the final manuscript.

