## Supplementary figure 1, figure 2, and table 1 for "Exposure of non-target white-footed mice (*Peromyscus leucopus*) to Second-generation Anticoagulant Rodenticides in an urban context"

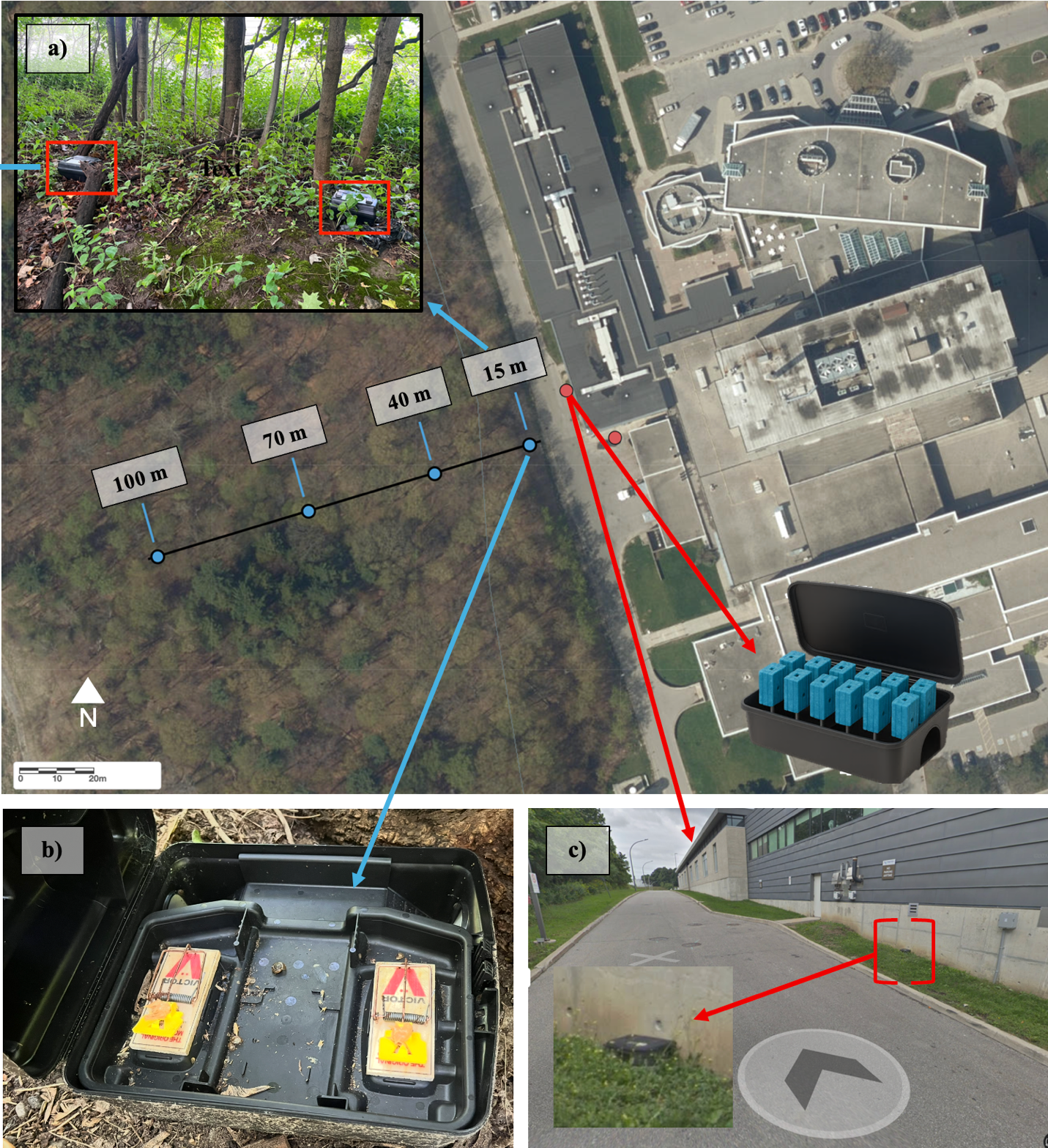
**Supplementary Material 1.** Trapping locations and transects: a) trap station placement and b) interior setup in relation to c) rodenticide bait stations at usage buildings adjacent to trap sites. Bait station image (above, right) generated with Chat GPT.

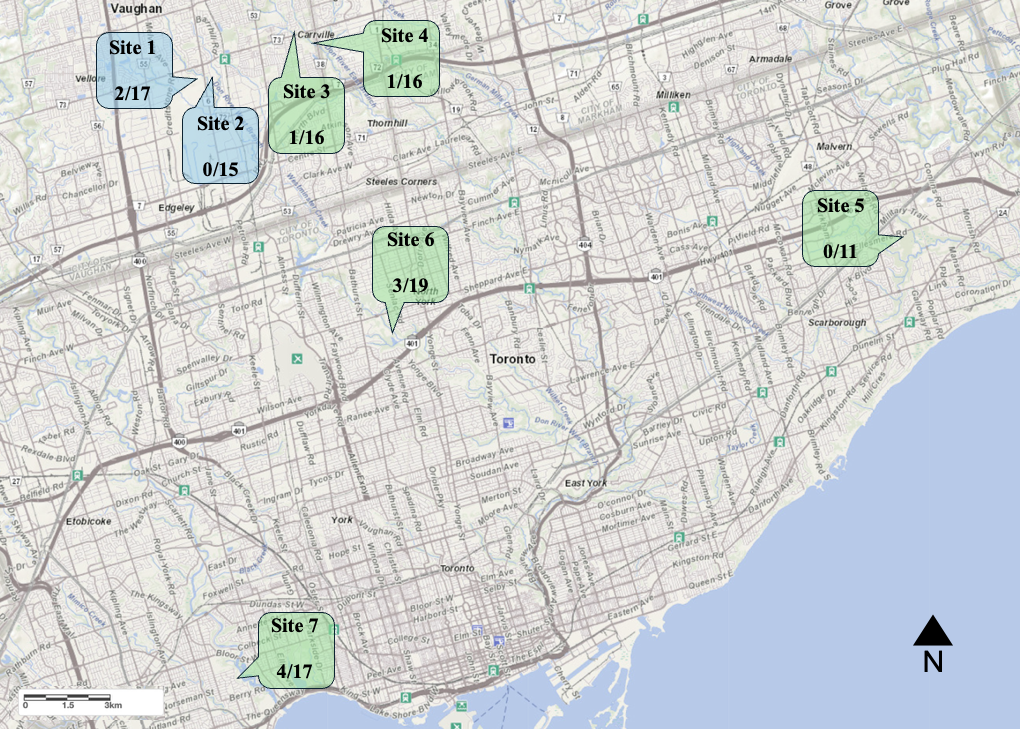

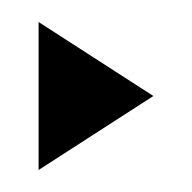

N

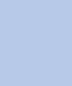

**Supplementary Material 2.** All seven sites in Toronto (four) and Vaughan (two) where trapped samples were tested for AR residues. Positives given as a fraction.

| Fisher’s Tests (exposed/unexposed) | | Logistic Regression | Linear Regression + PCA (exposure conc. ppm) | |
| --- | --- | --- | --- | --- |
|  |  | # bait stations ~ exposed/unexposed  p = 0.16 |  |  |
|  |  |  | Urban land (%) | p = 0.46 |
| Sex (M vs F) | p = 1 |  | Sex (M/F) | p = 0.35 |
| Ages (Imm. Vs Mat.) | p = 0.68 |  | Age (Mat./Imm.) | p = 0.37 |
| Sites (x7) | p = 0.44 |  | Julian Date of Capture | p = 0.57 |
|  |  |  | Distance (m) | p = 0.9 |
|  |  |  | PC1 | p = 0.66 |
| n = 111 | | n = 111 | n = 9 | |

**Supplementary Material 3** Comparison of exposure rates between groups (column one), exposure status (column two), and exposure concentration (ppm, column three) for those exposed to bromadiolone. No differences were observed, and no predictors were significant.
